# Metabarcoding of unfractionated water samples relates phyto-, zoo- and bacterioplankton dynamics and reveals a single-taxon bacterial bloom

**DOI:** 10.1101/058628

**Authors:** Christian Wurzbacher, Katrin Attermeyer, Marie Therese Kettner, Clara Flintrop, Norman Warthmann, Sabine Hilt, Hans-Peter Grossart, Michael T. Monaghan

## Abstract

Most studies of aquatic plankton focus on either macroscopic or microbial communities, and on either eukaryotes or prokaryotes. This separation is primarily for methodological reasons, but can overlook potential interactions among groups. We tested whether DNA-metabarcoding of unfractionated water samples with universal primers could be used to qualitatively and quantitatively study the temporal dynamics of the total plankton community in a shallow temperate lake. We found significant changes in the relative proportions of normalized sequence reads of eukaryotic and prokaryotic plankton communities over a three-month period in spring. Patterns followed the same trend as plankton estimates using traditional microscopic methods. We characterized the bloom of a conditionally rare bacterial taxon belonging to *Arcicella*, which rapidly came to dominate the whole lake ecosystem and would have remained unnoticed without metabarcoding. Our data demonstrate the potential of universal DNA-metabarcoding applied to unfractionated samples for providing a more holistic view of plankton communities.

## Introduction

Microbial communities are an integral component of total biodiversity (Barberán *et al.*, 2014) and play key roles in all ecosystems. Understanding of their composition and dynamics is a critical component of studying ecosystem functions and services. Plankton communities in freshwater and marine ecosystems are comprised of both microbial and macroscopic organisms from all three domains of life (archaea, prokaryotes, and eukaryotes). Traditionally, plankton is classified into functional groups such as phytoplankton, zooplankton, and bacterioplankton; or into size classes such as picoplankton, nanoplankton, and microplankton. This classification has resulted in the emergence of independent fields of inquiry, particularly the separation of prokaryotic and eukaryotic groups.

A consequence of this separation is that studies rarely survey all members of the plankton community simultaneously, except for a few contemporary marine surveys (Steele *et al.*, 2011, Lima-Mendez *et al.*, 2015). This is despite the potential that integrated studies have for providing an interdisciplinary view of plankton communities (Fuhrman *et al.*, 2015) by shedding light on the strength of biotic interactions (Needham and Fuhrman 2016). Most plankton studies employ size-selection steps (i.e. size fractionation by selective filtration) and genetic markers targeting either bacteria, archaea, or eukaryotes. Our literature review found that less than 0.5% of studies targeted all three (SI1). This tradition impairs a full integration of microbial communities into ecological concepts.

We used a universal 16S/18S primer pair to perform DNA-metabarcoding of unfractionated water samples in a study of the entire plankton community of a eutrophic, shallow, temperate lake (*Kleiner Gollinsee*) in northeastern Germany. We extracted total DNA from direct-filtered (0.2 μm) lake water (0.5 to 1 L), enabling us to screen all organisms from what is traditionally size-classified as pico- to approximately mesoplankton (per definition 0.2 μm - 20 mm). Our aim was to characterize the whole plankton community and its temporal dynamics in relation to algal biomass over a three-month period in spring (April – June) 2010. This is the period with the largest changes in plankton abundance and a high species turnover in most temperate eutrophic lakes. Our sampling (see SI1 for parameters and experimental procedures) was part of a larger, more traditional whole-lake survey of bacteria, phytoplankton and zooplankton from April 2010 to December 2011 (Brothers *et al.*, 2013, Hilt *et al.*, 2010) that we used for comparison.

## Results & Discussion

### Prokaryotic- and eukaryotic population dynamics

The DNA-metabarcoding of unfractionated water samples successfully amplified organisms across all three domains of life, yielding a total of 1986 bacterial, 544 eukaryotic, and 315 archaeal operational taxonomic units (OTUs) in the dataset. We recovered dominant organisms from nano- to mesoplankton size classes (see SI2 for taxa lists) including typical freshwater bacteria (e.g. *Polynucleobacter, Candidatus Aquirestis*), phytoplankton (e.g. *Cryptomonas, Synechococcus*), and zooplankton (e.g., *Cryptocaryon,* Diaptomidae). One field sample (June littoral zone sample) contained a small fish larva (inadvertantly sampled) and this was detected by DNA-metabarcoding as 1% of sequencing reads in that sample (classified as Cyprinidae, depicted as Teleostei in SI2). Archaeal sequences were not abundant in our Lake Gollin samples. This was not surprising because the lake was oxic during the sampling period and Archaea are rarely found in oxygenated freshwaters (Pernthaler *et al.*, 1998, Gies *et al.*, 2014). The low number of archaea was not likely caused by a primer bias, because the primer pairs have been used successfully to detect a dominance of *Archaea* in the anoxic zone of a meromictic lake (Gies *et al.*, 2014). We observed a pronounced shift in relative read abundance from a dominance of eukaryotes in April to a dominance of prokaryotes in June for all sampled water compartments (i.e. littoral, pelagic and sediment zones; Fig. 1a). This was accompanied by an increasing heterotrophs:phototrophs ratio (SI3).

**Figure 1.**
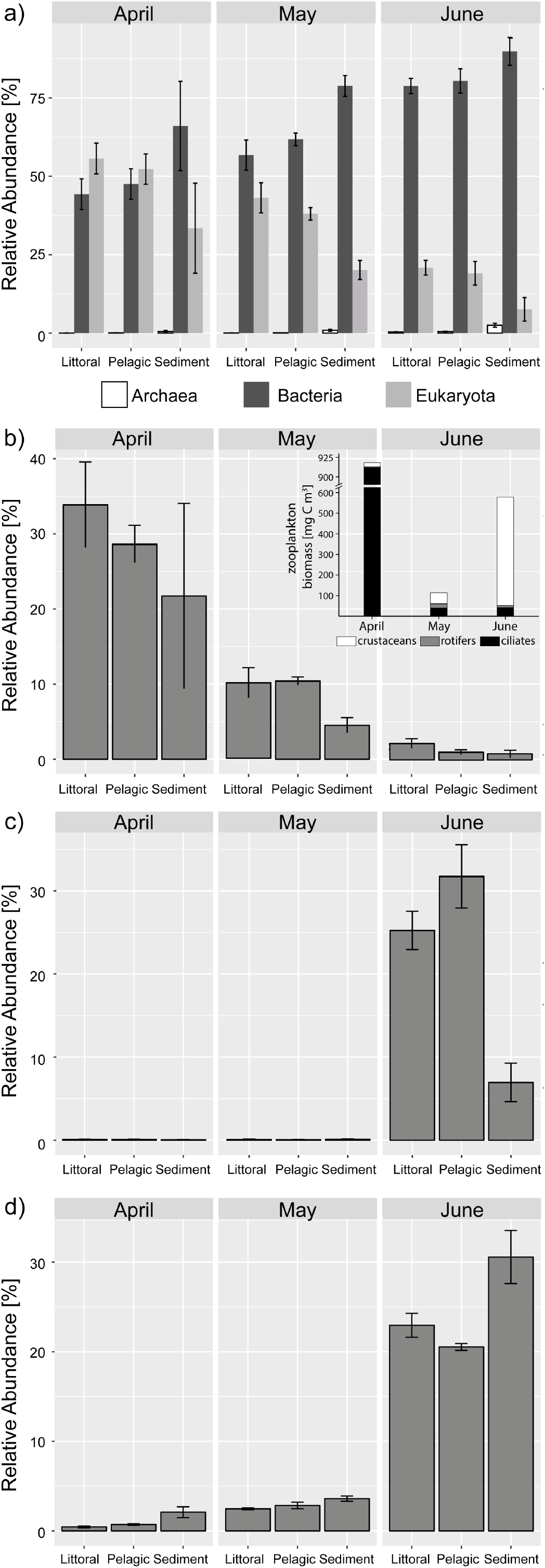
Spatial and temporal changes of microorganisms in Lake Gollin from three locations (littoral, pelagic, and above sediment; sampled as triplicates) per sampling event (21. April 2010, 19. May 2010 and 17. June 2010). The panels display sequence abundance (mean ± standard deviation) based on proportions of (a) all three domains with a total of 1986 bacterial, 544 eukaryal, and 315 archaeal OTUs in the dataset; (b) the sum of all ciliate OTUs (89 OTUs) with zooplankton biomasses in the same month obtained with traditional microscopic counting in the right hand corner, (c) the dominant *Arcicella* OTU, and (d) the dominant *Variovorax* OTU. Amplicons were based on the V9 region of the ribosomal small subunit (SSU: 16S/18S) for taxa detection (Engelbrektson *et al.*, 2010, Gies *et al.*, 2014). Methodological discussions on a related cross-domain single marker can be found in Parada *et al.* (2016). Sequencing followed the procedures described by Hölker *et al.* (2015) with the modification that we employed the AccuPrime High Fidelity Polymerase (Invitrogen, Carlsbad, USA). Sequences were processed in Mothur (version 1.24.1; Schloss *et al.*, 2009) and classified with SINA aligner (version 1.2.11; Pruesse *et al.*, 2012) against the SILVA SSU reference database (115 Ref NR 99, www.arb-silva.de). For details on experimental procedures see SI1.

### Comparison with microscopical observations

Abundance patterns based on DNA-metabarcoding data followed the trend of the sum parameters of phyto-, zoo- and bacterioplankton obtained from traditional microscopical counting data (e.g. ciliates in Fig. 1b, SI3). We recovered all microscopically counted planktonic organisms, although there were frequent mismatches between the classification depths in the two methods (Table S2 in SI1). Counting and sequence data were not taken on the same day and are therefore not directly comparable. Nonetheless, accounting for this difference by averaging, we found a rank-based correlation between phytoplankton reads and phytoplankton counts (Spearman's rho = 0.66, p < 0.001). For zooplankton this relationship was not significant (p > 0.05), although subsets exhibited a strong correlation (e.g. certain groupings of ciliates). The strongest correlation between sequence and count datasets was found with log-log transformed data for both phytoplankton (Pearson's r = 0.45, p < 0.001) and zooplankton (r = 0.37, p < 0.05).

Our data also captured the temporal dynamics in reported species abundances of the lake. A significant fish-kill event occurred in the winter prior to our study, caused by a prolonged ice-cover that led to anoxia (Hilt *et al.*, 2015). This had an important impact on the lake ecosystem, and a bloom of herbivorous ciliates in April 2010 (Lischke *et al.*, 2016) was clearly visible in our sequence data (approx. 30% of all reads, Fig. 1b). The ciliates would have exerted strong grazing pressure on the small plankton (< 5 μm; Lischke *et al.*, 2016) but the ciliate population crashed in May-June (Fig. 1b) and the role of grazer was then filled by crustaceans (Hilt *et al.*, 2015). Similarly, our sequence data from the pelagic zone indicated shifting crustacean:rotifer:ciliate read ratios, being 0:6:56 in April, 9:1:26 in May, and 38:6:5 in June (SI3). The replacement of ciliates by crustaceans may have opened a niche for the observed bacterial dominance in June, via reduced grazing pressure and increased substrate supply via sloppy feeding of the copepods.

### Bloom-forming OTUs

The bacterial dominance occurred in June and was attributed to OTUs classified as *Arcicella* and *Variovorax* (Fig. 1c,d). June was also the period in which the highest bacterial carbon production was measured (SI3, Brothers *et al.*, 2013; Lischke *et al.*, 2016). A single *Arcicella* OTU was most abundant in the pelagic open water and its greatest proportion of reads (40%) was observed at 1 m sampling depth (Fig. 1c), suggesting it colonized the water surface. *Variovorax* was more prevalent above the sediment, suggesting colonization from the sediment (Fig. 1d). In contrast to *Variovorax*, which exhibited already stable abundances at the other two sampling dates, *Arcicella* was present at very low abundances in April and May (<0.2%) and can thus be classified as a conditionally rare taxon (Lynch and Neufeld, 2015). There are few reports of blooms of rare bacterial taxa correlated with algal blooms. Gilbert *et al.*, 2012 described a *Vibrio* sp. bloom in the English channel and Bizic-Ionescu *et al.* (2014) described the genera *Flavobacterium* and *Undibacterium* associated with a phytoplankton breakdown event in a lake. In order to test whether *Arcicella* was a reoccurring taxon in Lake Gollin or if this was a unique appearance related to the fish-kill disturbance, we screened additional samples (two size-fractions in this case: 0.2-5 μm & > 5 μm, from 19 monthly taken samples) that were available from Lake Gollin (Brothers *et al.*, 2013). A*rcicella* occurred again in the following year (Fig. 2) and its periodical appearance may be negatively related to chlorophyll-a concentrations and positively to crustacean biomass (Fig. 2b; see Table S3 in SI1 for correlations). This bacterial bloom therefore appears to be different from previously described blooms which were positively linked to phytoplankton abundance (e.g. Gilbert *et al.*, 2012). We conclude that it was likely related to the fish-kill (see above), and our data on microbial species turnover also suggested opening of a niche allowing for bloom development of a single bacterial species. Conditionally rare taxa can thus have a disproportional effect on community structure (Shade *et al.*, 2014), although knowledge of their ecological and metabolic potential is required better understand their ecosystem-wide consequences.

**Figure 2.**
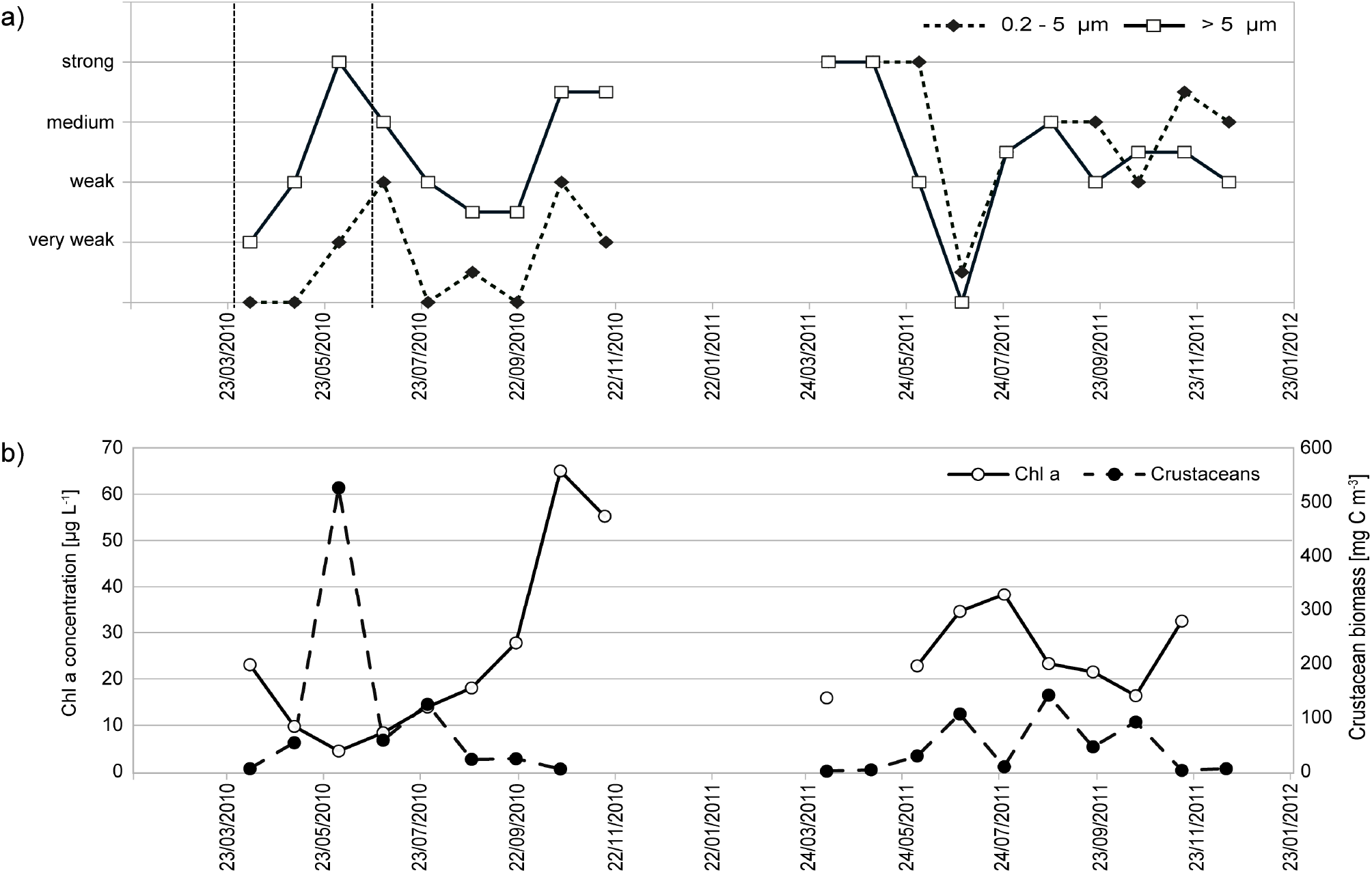
Seasonal appearance of (a) *Arcicella* exhibited pronounced maxima and minima over the course of the 2 years and appeared in the particle-attached (> 5 μm) and free-living fraction (0.2 - 5 μm). *Arcicella* was detected using a PCR assay (see experimental procedures S1) and evaluated based on gel electrophoresis band intensity where 0 = no PCR product, 1 = very weak product, 2 = weak product, 3 = medium product, 4 = strong product. For this data we applied rank based correlation tests based on local similarities for time-series analysis (see Table S3 in SI1). Correlations that were significant correlated with the band intensity of Arcicella products derived from both size fractions were Chlorophyll a and crustacean biomass, both of which were plotted in panel (b).

### What is Arcicella?

We searched the freshwater and marine literature and found few reports of *Arcicella* in environmental samples. *Arcicella* was the second-most abundant OTU (6.8%) in a study of 5 large arctic rivers (Crump *et al.*, 2009), although the taxon is not further discussed in their article. In some cases, *Arcicella* was mentioned in a figure legend or in supplementary materials and we had to request abundance data from the authors. One *Arcicella* OTU was among the most dominant OTUs in the Danube river although it comprised only 2% (± 2%) of relative read abundance (D. Savio pers. comm.; Savio *et al.*, 2015). A single *Arcicella* OTU comprised 12 (± 4%) of the relative read abundance in a small turbid glacial lake (Peter pers. comm.; Peter and Sommaruga, 2016). In spite of these observations, their autecology has never been investigated.

### Study limitations and perspectives

There was a large standard deviation among replicates for crustaceans and rotifers (depicted as zooplankton in S3). These organisms belong to the mesoplankton size class (0.2 to 20 mm) and it may be necessary to increase the water volume of samples to better quantify them. Sequences belonging to Metazoa could only be classified to phylum-class level (e.g. Maxillopoda, Rotifera, Teleostei) using the automatic classifier. One reason for this is that the SILVA reference taxonomy that we used has not implemented freshwater zooplankton yet, but we partially resolved this problem by using the original taxonomy provided by NCBI/EMBL. A second reason is that the SSU region targeted here does not provide sufficiently variation for many Metazoan groups (Tang *et al.*, 2012). We therefore estimated the resolution of the universal marker for all of the planktonic organisms in Lake Gollin (see Table S2 in SI1). Most Rotifera OTUs could be classified only to order, while most Maxillopoda OTUs could be classified to family or genus level. Differing taxonomic resolution among groups (in particular Metazoan taxa) is one limitation of using a universal marker, and this should be considered in future studies of these groups. The effect is most obvious when employing a fixed OTU cut-off such as 97%. One solution could be to combine universal metabarcoding with a dynamic cutoff method focused on noise removal than on clustering (e.g., Eren *et al.*, 2015). Such an approach may increase the phylogenetic resolution. Moreover, biases in relative taxa abundance can be introduced through DNA extraction, PCR (primer choice, amplification), library preparation and sequencing steps (Gilbert *et al.*, 2012; Singer *et al.*, 2016). As a result, quantitative estimates are likely to be semi-quantitative at best. Universal metabarcoding has the same limitation but at least comes with the advantage, that it provides a balance between all organisms, since most of the DNA template will be derived from the target groups (only excluding viruses in this case). To date, it has produced conclusive results for the general trends in plankton communities (this study, see also Gies *et al.*, 2014, Parada *et al.*, 2016, Needham and Fuhrman, 2016).

## Conclusions

Using unfractionated water samples with universal DNA metabarcoding allowed us to document major changes in almost the entire size- and functional spectrum of freshwater plankton with a single water sample analysis. Changes in the relative abundance of OTUs closely matched the seasonal dynamics of phyto-, zoo-, and bacterioplankton reported for this lake based on microscopy, indicating that relative abundance data based on read counts are ecologically meaningful. The discovery of a bloom of a largely overlooked freshwater bacteria genus was remarkable, with potential implications for the whole lake ecosystem. Our results highlight the potential of simultaneously studying both microbial and macrobial communities for an improved understanding of whole ecosystem changes. Integrative and interdisciplinary analyses may help to answer broad ecological questions in freshwater systems related to the role of keystone species; ecosystem resilience and resistance; and cross-domain interactions of species. Unfractionated sampling coupled with metabarcoding using a universal primer provides a powerful approach for studying plankton dynamics in aquatic systems and shows promise for long-term whole community monitoring.

## Acknowledgements

We thank Uta Mallock for chemical analysis, Susan Mbedi for assistance in sequencing and Marie Jeschek for assistance in data processing. Financial support was provided by the SAW/Pact for Research and Innovation of the Leibniz Association projects “Terralac”, “TEMBI”, and “MycoLink”. R. Tischbier (Stiftung Pro Artenvielfalt) kindly provided us access to the lake. The authors declare no conflict of interest.

